# Nonhomologous tails direct heteroduplex rejection and mismatch correction during single-strand annealing in *Saccharomyces cerevisisae*

**DOI:** 10.1101/2022.11.15.516560

**Authors:** Elena Sapède, Neal Sugawara, Randall G. Tyers, Yuko Nakajima, Mosammat Faria Afreen, Jesselin Romero Escobar, James E. Haber

## Abstract

Single-strand annealing (SSA) is initiated when a double strand break (DSB) occurs between two flanking repeated sequences, resulting in a deletion that leaves a single copy of the repeat. We studied budding yeast strains carrying two 200-bp *URA3* sequences separated by 2.3-kb of phage lambda DNA in which a site-specific DSB can be created by HO or Cas9 endonucleases. Repeat-mediated deletion requires removal of long 3’-ended single-stranded tails (flaps) by Rad1-Rad10 with the assistance of Msh2-Msh3 and Slx4. A natural 3% divergence between repeats (designated F and A) causes a significant reduction in the frequency of SSA repair. This decrease is caused by heteroduplex rejection in which mismatches (MMs) in the annealed intermediate are recognized by the MutSα (Msh2 and Msh6) components of the MM repair (MMR) pathway coupled to unwinding of the duplex by the Sgs1-Rmi1-Top3 helicase. MutL homologs, Mlh1-Pms1 (MutLα) are not required for rejection but play their expected role in mismatch correction. Remarkably, heteroduplex rejection is very low in strains where the identical repeats were immediately adjacent (Tailless strains) and the DSB was induced by Cas9. These results suggest that the presence of nonhomologous tails strongly stimulates heteroduplex rejection in SSA. DNA sequencing analysis of SSA products from the FA Tailed strain, where the F variant carries six single-base mutations and one +1(T) insertion within a run of 10 Ts showed a gradient of correction favoring the sequence opposite each 3’ end of the annealed strand. Mismatches (MMs) located in the center of the repair intermediate were corrected by Msh2-Msh6 mediated mismatch correction, while correction of MMs at the extremity of the SSA intermediate often appears to use a different mechanism, using either 3’ nonhomologous tail removal or 3’ to 5’ “proofreading” resection by DNA polymerase δ followed by synthesis to fill in the gap. In contrast, in FA Tailless strains there was a uniform repair of the MMs across the repeat, with a bias in favor of the “left” copy. A distinctive pattern of correction was found in the absence of *MSH2*, in both Tailed and Tailless strains, while the deletion of *MSH6* resulted in unrepaired MMs.

## INTRODUCTION

In the human genome, repetitive DNA sequences such as LINEs (long interspersed nuclear elements) and SINEs (short interspersed nuclear elements, such as Alu) constitute at least 34% of the genome [1]. Recombination between repetitive DNA sequences can lead to chromosomal translocations, deletions, or inversions which are thought to contribute to tumor formation [2]. As one example, many of the mutations in *BRCA1*, a gene known to be linked to breast and ovarian cancers are due to deletions and duplications between Alu repeats [3].

One way in which a double-strand break (DSB) can be repaired is through single-strand annealing (SSA) between repeated sequences flanking the DSB, resulting in a chromosomal deletion retaining one copy of the repeated sequence, also referred to as repeat-mediated deletion (RMD). SSA requires extensive 5’ to 3’ resection of the DSB ends to expose complementary single-strand DNA (ssDNA) sequences that can anneal, producing an intermediate in which there are 3’-ended single-stranded nonhomologous tails. Both in yeast and in mammals, SSA can occur between repeats that are as far apart as 25 kb [4, 5]. Extensive resection in yeast depends on either the Exo1 or Sgs1-Rmi1-Top3-Dna2 exonucleases; in mammals the Sgs1 homolog BLM appears to play a similarly important role in resection [6]. Strand annealing in budding yeast requires the Rad52 annealing protein but is independent of the Rad51 recombinase that is required for most other types of homologous recombination (McDonald and Rothstein; Ivanov et al. 1996). In mammals, Rad52 appears to be one of several annealing proteins [6]. In yeast, completion of SSA requires the excision of nonhomologous tails by the Rad1-Rad10 (ERCC1-XPF) 3’ flap endonuclease and the Slx4 scaffold protein [7, 8]. In yeast, efficient flap removal is also dependent on the MutSß Msh2-Msh3 mismatch repair proteins that recognize branched DNA structures [9–11].

The distance between the DSB and the repeated fragments defines the amount of end resection required for SSA repair. In *S. cerevisiae*, a DSB created between repeats as far apart as 25 kb led to the efficient deletion of the separating fragment [5]. Mutations such as *fun30*Δ and *rad50*Δ that impair resection prevent completion of SSA when the repeats are separated by a long distance [12]. In mammals, studies on mouse embryonic stem cells showed that end resection is critical for SSA at least within the first few kb (3.3 kb) [4].

Besides the distance between the repeated fragments, another parameter that influences the efficiency of SSA is the degree of homology between the repeats. In both yeast and mammals, a low level of heterology - as little as 3% - decreased SSA repair more than two-fold [4, 10]. In budding yeast, this reduction is caused by heteroduplex rejection, in which the annealed resected strands of the repeat are unwound by the Sgs1-Rmi1-Top3 helicase complex, apparently prompted by the binding of the Mutsα Msh2-Msh6 mismatch repair proteins, but independent of the MutLα Mlh1-Pms1 mismatch repair proteins that are still required for mismatch correction [10, 13]. Heteroduplex rejection does not involve the degradation of the annealing sequences, as they can participate in other, more distant SSA events with a fully homologous partner [10, 11].

The relationship between heteroduplex rejection and the removal of a 3’-ended nonhomologous tail has been unclear. In a Rad51-dependent recombination process such as break-induced replication (BIR), the presence of 1 or 2 mismatches between 108-bp regions undergoing recombination had little or no effect when the DSB end was perfectly aligned with the donor, but these same heterologies reduced BIR substantially when the DSB end carried a 68-nt nonhomologous tail [14]; even a 3-nt tail had a significant effect. Thus, we wished to assess how nonhomologous tails affected heteroduplex rejection in SSA.

We used a simple SSA assay [10] in which a site-specific DSB is induced between two 200-bp regions located in the 5’ promoter region of the *URA3* gene on chromosome V. These duplicated regions were derived from two different *S. cerevisiae* strains, designated F and A, which harbor six single base substitutions and a T insertion in F relative to A (Figure 1A); these naturally occurring heterologies are nonrandomly spaced as shown in Figure 1B. The insertion/deletion of T is found in a run of 12/10 Ts immediately adjacent to mismatch 3 (MM3).

**Figure 1.**
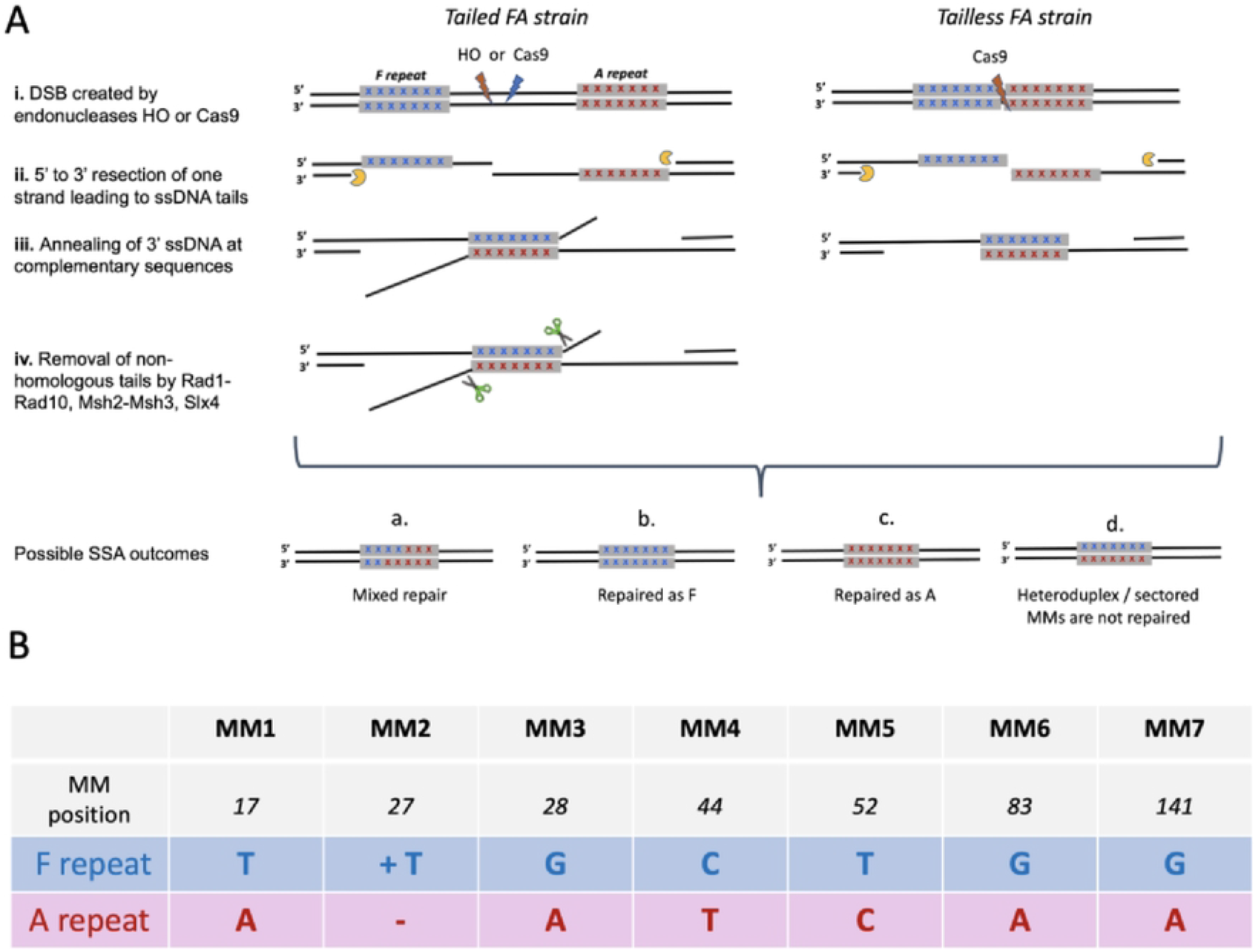
SSA repair in *S. cerevisiae.* A. SSA repair between divergent repeats. (i) A DSB is created by HO or Cas9 endonucleases between two 200-bp repeated sequences in Tailed (left) and in Tailless (right) strains. In FA Tailed strains the repeated fragments are separated by a 2.6 kb DNA sequence. (ii) DSB ends undergo 5’ to 3’ resection, creating 3’ single-strand DNA tails. (iii) Annealing of single-strand DNA at complementary sequences creates an intermediate with mismatched base pairs from which (iv) the nonhomologous tails are removed by Rad1-Rad10 endonuclease with the assistance of Msh2–Msh3. Depending on how MMs are corrected, there are different possible outcomes of SSA: Mixed repair (a); all MM are repaired as F (b); all MM are repaired as A (c); If mismatches are not corrected, progeny containing both genotypes will result in a sectored colony (d). B. Location of MMs. Nucleotides at MM positions are shown for the F fragment (in blue) and from the A fragment (in red).

In the FA Tailed strain, the F and A repeats are separated by 2.3 kb segment of phage λ DNA into which is embedded a 117-bp recognition site for the site-specific HO endonuclease [10, 11] (Figure 1A). DSBs can also be created at different locations in the phage λ DNA through the use of CRISPR/Cas9 (Figure S1.A). We also gene-edited this construct to place the F and A segments immediately adjacent to each other (see Materials and Methods) such that the junction can be cleaved precisely by Cas9 (Figure 1A & Supplementary Table 1), without creating a nonhomologous tail (termed as Tailless strain).

We show that heteroduplex rejection strongly depends on the presence of the 3’-ended nonhomologous tails. The lack of nonhomologous tails led to more efficient SSA in both FA and AA Tailless strains compared to Tailed strains. We show further that the pattern of mismatch repair changes dramatically in the absence of nonhomologous tails. When tails must be clipped off to complete SSA, mismatch correction favors retention of the allele on the opposite strand, producing a gradient of repair from the two ends. In contrast, in the absence of tails, assimilation of each mismatch occurred coordinately across the entire sequence. In this process a critical role is played by Msh2, which has multiple functions during SSA repair, including heteroduplex rejection, nonhomologous tail removal and mismatch correction.

## MATERIALS AND METHODS

### Strains

Strain tNS1379 carries a duplication of 200-bp sequences upstream of the *URA3* open reading frame (designated AA) separated by 178 bp of pUC9 DNA, the HO endonuclease cut site (117 bp) derived from *MAT*a [15], and 2.3-kb phage λ DNA. In tNS1357 (FA Tailed strain) the leftward *URA3* repeat was replaced by sequences derived from strain FL100 and differs from the AA Tailed strain by six single-nucleotide substitutions and one indel within a run of 12 Ts [10] (Figure 1.B & Supplementary Table 1 & Supplementary Table 2).

Tailless strains were obtained by gene editing tNS1379 (AA Tailed strain) and tNS1357 (FA Tailed strain). A DSB was created by Cas9 cleavage between the two repeated fragments and a 100-bp dsDNA designed to have 50 bp homology to the 3’ and 5’ ends of the left and right repeated fragments was used as a template for repair, allowing the removal of 2.3kb phage λ DNA and other intervening sequences, resulting in a strain with two immediately adjacent repeats, designated as FA and AA Tailless strains (Supplementary Table 1, 2, 3 & 4).

Additional gene editing was employed to obtain an AF Tailed strain (where the positions of the A and F segments are reversed), nFA or AnF Tailed strains (when the insertion of 1 T from F fragment was replaced by a G) and FA Tailed strains with PAMs adjacent to each repeated fragment. DSBs were created by: pES6 (an Leu2-marked Cas9 plasmid that cuts upstream of the F repeat – used to create nF strains); pES11 (HPH Cas9 plasmid that cuts downstream of the A fragment – used to create AF strain from the AA strain); pES8 (HPH-marked Cas9 plasmid that cuts upstream of the A repeat was used to insert a PAM adjacent to the right fragment), and pES18 *(LEU2-marked* Cas9 plasmid that cuts downstream of the F repeat, was used to insert a PAM adjacent to the left repeat), in each case using an single-stranded oligonucleotide to edit the site. The dsDNA templates used for editing were obtained by PCR amplification and the ssDNA templates were ordered from IDT (Supplementary Table 4).

Deletions of *SGS1, MSH2, MSH6, MSH3, RAD51, RAD52, MLH1*, and *PMS1* were performed by replacement with KANMX, NATMX, or HPHMX cassettes using PCR mixed primers with 40-bp homology to sequences 5’ and 3’ to the open reading frame and 20-bp homology to MX cassettes [16]. Tailed and Tailless strains were transformed with the MX cassettes using high efficiency yeast transformation methods [17].

### Analysis of SSA

HO endonuclease under the control of a *GAL<u>1</u>,10* promoter (*GAL::HO*), carried on a centromeric plasmid, pFH800 [18], was induced by addition of 2% galactose to create a single DSB between the 200-bp repeated segments [10]. After induction of *GAL::HO* nearly all survivors were SSA products and could thus be monitored by colony counting. Cells were plated for individual colonies on YEPD and on YEP-GAL plates to induce HO endonuclease. After 2 days, cells that grew on YEPD and on YEP-Gal plates (DNA break survivors) were replica plated on tryptophan drop-out plates (the marker for pFH800 plasmid) to determine SSA frequencies. Approximately 10% of cells lost the *TRP1*-containing plasmid. SSA repair was confirmed by PCR using primers NS154-pUC and URA3p14 upstream and downstream from the two repeated fragments (Supplementary Table 4) a 250 bp fragment indicates SSA repair; in the absence of DSB the PCR product is 2.9 kb.

### Galactose-inducible Cas9 Endonuclease pRTO2

A gRNA targeting sequence designed to target a region between the adjacent repeats in Tailless strains was cloned into a *LEU2*-marked plasmid (bRA77), in which Cas9 is expressed under the *GAL<u>1</u>,10* galactose-inducible promoter [19]. Tailless strains were then transformed with Cas9 plasmids by using the conventional LiAc protocol as previously described [19]. To induce the Cas9 endonucleases activity, cells were grown for 3 h in YP-lactate, and then ~500 cells were plated on YEPD and YEP-Gal plates. After 2 days of incubation at 30°C, colonies were replica-plated to plates lacking leucine to score cells that retained the Cas9 plasmid, after which SSA viability was scored.

### Constitutive Cas9 plasmids

pAB101, an HPH Cas9 plasmid targeting a site in the 117-bp HO cleavage site from *MAT***a** adjacent to the phage λ sequence [10] was used to create a DSB 268 bp downstream Left (F) Fragment. This Cas9 cleaves adjacent to the site cut by HO endonuclease (Figure S1A). gRNA sequences are listed in Supplementary Table 5; pES02 is a HPH-marked Cas9 plasmid that cuts approximately in the middle between the two repeated fragments, while pES08 is an HPH-marked Cas9 plasmid that cuts 24 bp upstream of the right (A) fragment (Figure S1A). To create DSBs with only one nonhomologous tail we designed gRNAs to direct Cas9 cutting adjacent to the F fragment (pES19 – nonhomologous tail on bottom strand) or the A fragment (pES20 - nonhomologous tail on the top strand) Figure 2C. To investigate SSA frequency when DSBs are created at different locations on λ DNA (on 2.3-kb tail), the FA Tailed strain was transformed with one of the plasmids (pAB101, pES2, pES8, pES19 and pES20) in the presence of a second, *LEU2*-marked plasmid to normalize for variations in transformation. bRA89 is a HPH-marked plasmid lacking Cas9 that was used as a control. Transformants were plated directly on *LEU2* drop-out plates (marker of the normalization plasmid) and replica-plated to HPH plates after three days of incubation at 30°C. The survivors carrying both *LEU2* and HPH plasmids were counted. The number of colonies transformed with pAB101, pES2, pES8, pES19 or pES20 (DSB repaired colonies) were compared to the number of colonies transformed with bRA89 (no DSB).

**Figure 2.**
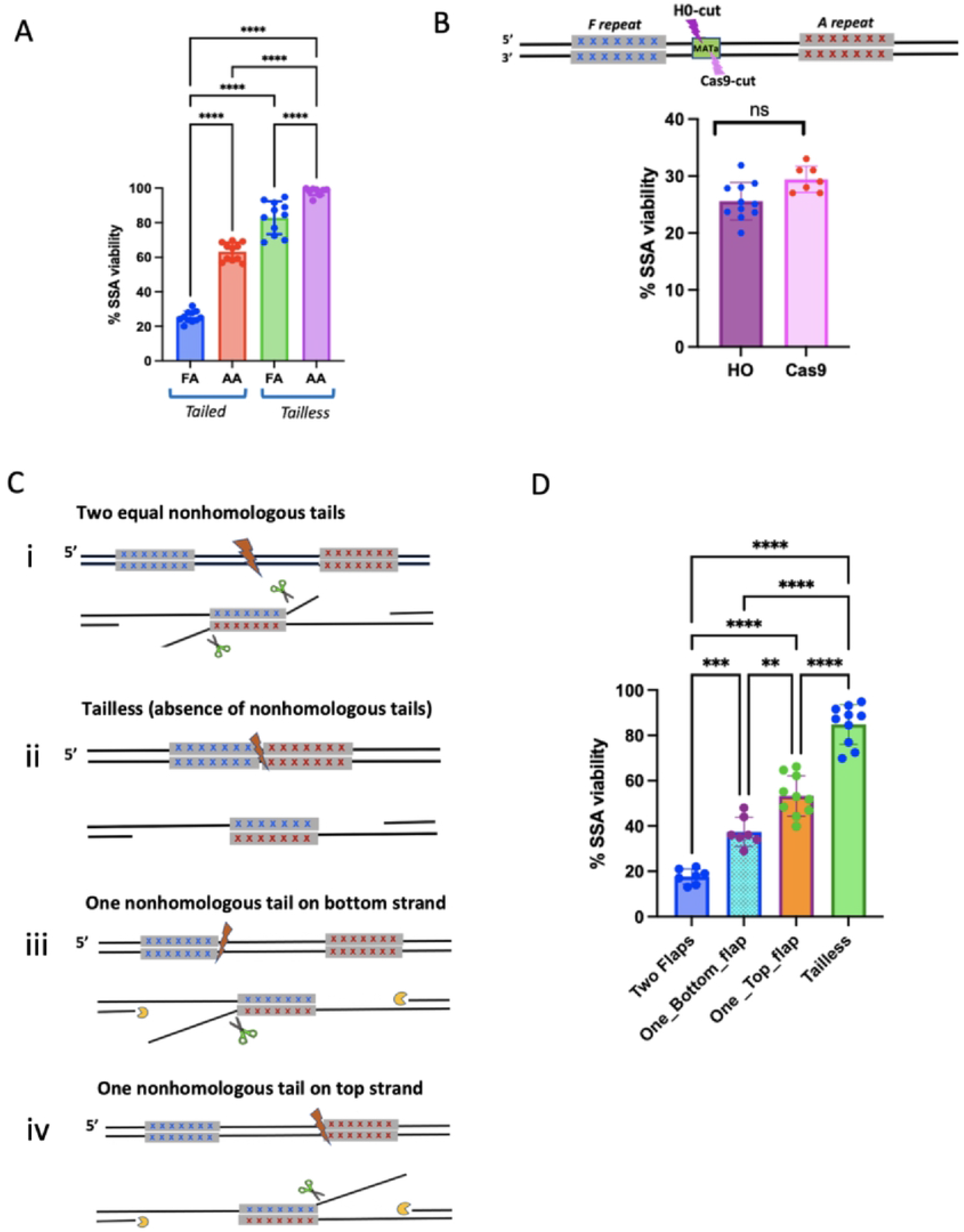
SSA repair frequency A. Both mismatches and nonhomologous tails reduce SSA. The frequency of DSB-induced SSA was measured by after induction of *GAL::HO* or *GAL::CAS9* compared to cells grown on dextrose. Statistical test: One-Way ANOVA; Error bars represent the standard deviations; **** = p< 0.0001. B. DSB Comparation: HO vs Cas9 endonucleases - Viability in FA Tailed strains where the DSB was created by HO or by a Cas9 (pAB101) endonuclease targeting the same locus (MATa). Experiments were independently repeated at least five times. Error bars indicate standard deviations. Experimental means were statistically compared by the unpaired t-Test. C. SSA intermediates when a Cas9-mediated DSB is created (i) midway between the divergent fragments; (ii) in Tailless strains; (iii) adjacent to the left fragment resulting in a 2.4 kb nonhomologous tail (flap) on the bottom strand, 17 bp upstream of MM1 or (iv) adjacent to the right fragment resulting in a 2.7 kb nonhomologous tail (flap), on the top strand, 60 bp downstream MM7. D. Effect of nonhomologous tails on SSA: FA tailed strain was transformed with pES02 (creating two equal nonhomologous tails) or with pES19 and pES20 to create a single flap on the bottom strand or top strand, respectively and compared after galactose induction. Experiments were independently repeated at least than five times. Experimental means were statistically compared by One-Way ANOVA test. Error bars represent the standard deviation (SD) of n > 5 viability assays; ** p = <0.01; *** p < 0.001; **** p< 0.0001

Experiments were independently repeated 5 to 9 times. Experimental means were statistically compared by a one-way ANOVA test.

### Analysis of mismatch repair after SSA

Genomic DNA was obtained from single colonies that were boiled in 50 μL of Lysis Buffer (10mM Tris-HCl, 2mM EDTA, 1% SDS). The 250-bp products confirming SSA repair were PCR-amplified using primers (NS154-pUC as forward and URA3p14 as reverse primers) that were positioned 50 bp outside the repeated fragments. The amplification was performed in 20 μL of reaction (2XPCR FailSafe^TM^ PreMix E from Biosearch Technologies, 1.25 U of GoTaq DNA Polymerase (Promega), < 1μg DNA template and 0.5 μM of each specific primers. The amplification consisted of an initial activation step at 95 °C for 5 min followed by 25 cycles, including a denaturation step at 95 °C for 0.30 min, annealing at 55 °C for 0.30 min, and extension at 72 °C for 1 min. The final extension was performed for 5 min at 72 °C. Individual PCR reactions were sequenced (Genewiz) using the reverse primer URA3p14, and the traces were aligned with the A and F sequence fragments. The chromatogram of the SSA repair products could show either the nucleotide from the A fragment or F fragment or the presence of both A and F alleles, suggesting that the MM wasn’t repaired or that a colony derived from post-replicative cells in which repair occurred independently in two daughter nuclei. Approximately 50 samples were sequenced for each construct. Sequence analyses was carried out using the alignment feature of GENEIOUS 11.0.5 software.

### Strand linkage analysis – Barcode NGS

Sanger sequencing described above can show that one or more sites retained both F and A alleles, but cannot assign linkage of the markers. To determine which alleles were carried on a single DNA strand, we performed paired-end sequencing of individual SSA outcomes. Strand-specific analysis of SSA progeny was done as follows. We built six libraries: FA_WT Tailed, *FA_msh2*Δ Tailed, FA_*msh6*Δ Tailed, FA_WT Tailless, FA_*msh2*Δ Tailless, FA_*msh6*Δ Tailless. Each library contains sequences amplified from 96 SSA repaired colonies using 8 specific forward primers and 12 specific reverse primers (with 8-bp barcodes; Supplementary Table 4). Genomic DNA was obtained from single colonies as described above. Each independent SSA product was amplified for 25 cycles, using a unique pair of barcode primers in 20 μL of PCR mix using GoTAQ or Q5 polymerase. All 96 barcoded PCR products from each library were pooled, DNA purified, and analyzed using Amplicon-EZ; which uses Illumina Mi-seq technology (Azenta Life Sciences, Chelmsford, MA). The R1 and R2 paired-end reads were joined by BBMerge [20] at the highest stringency presetting. Length and quality score filtering (L>250 bp, Q>35, respectively) was done using FastP [21]. Mapping and quantification of sequence variants was done using the Variable Antigen Sequence Tracer (VAST) [22] with the F-sequence as the reference and the A-sequence as the cassette. VAST variant_frequency reports contain frequency information for all observed recombinants between the reference and cassette(s) organized by colony and genotype. TSV files containing this information were processed using FA_variant_processor_MM2_fix.py, a Python script that labels MM positions as being either from the reference (F-sequence) or cassette (A-sequence) and treats sequences where the second position (MM2) does not agree with the 1st (MM1) and 3rd (MM3) positions as errors, and corrects them. For example, the FAFFFFF sequence is counted as if it were FFFFFFF. Each colony is then assigned a duplex genotype by picking the two most frequent sequences that have a frequency greater than 19%. Details and code are available from: https://github.com/users/RandallTyers/FA_variant_processor.

### Rationale for MM2 error correction

Mismatch 2 (MM2) is a single-base insertion/deletion in a run of 10 Ts. Poly(dT:dA) stretches as short as T_12_ are notable for blocking DNA polymerases both in vitro and in vivo [23, 24]. To establish the fidelity of identifying the identity of MM2 in repaired SSA events we analyzed DNA sequences derived from strains that either carried only F or only A alleles (i.e. MM2 was either T_12_ or T_10_). Using DNA polymerase Q5 (High Fidelity DNA polymerase-M0491L from NEB) we found that 5 ± 2% of the reads appeared to be different from the template sequence in this region; such changes could come either from the PCR amplification or from subsequent steps in DNA sequencing. As has been previously reported [25] Taq polymerase generated single nucleotide indels at a much higher frequency (in our samples as high as 24±2%). We note that the position of the indel within the polyT sequence cannot be determined. In analyzing SSA events between F and A alleles, in those instances when both MM1 and MM3 were derived from the same template, we assumed that MM2 was also retained/corrected in the same direction. The fraction of cases in which a given mismatch was changed to that of its neighbor is shown in Figure S8.

## RESULTS

### Nonhomologous tails stimulate heteroduplex rejection

We used a viability assay to evaluate the difference between Tailed and Tailless SSA strains after creating a DSB between the repeated fragments (Figure 1A). When the repeats were identical, viability was 68% in the AA Tailed strain and 100% in AA Tailless strains (Figure 2A), suggesting that the presence of nonhomologous tails derived from sequences between the repeated sequences inhibits repair. In FA stains, The 3% divergence between the repeats led to a decrease in repair frequency to only 26% in FA Tailed strains, 2.5-fold lower than the equivalent AA Tailed strain, whereas in the FA Tailless strain the viability was 85%. Hence there is some impact on viability imposed by the seven mismatches, but this is more pronounced in the presence of 3’ nonhomologous tails that must be removed to complete SSA. Together, these results suggest that both the presence of nonhomologous tails and the mismatches (MMs) lead to heteroduplex rejection that causes a deficit in repair (Figure 2A). Analogous results were obtained when Cas9 encoded in plasmid pAB101 was used to cleave a site close to the HO cleavage site (Figure 2.B).

### Effect of DSB position between repeated sequences

To investigate how only one nonhomologous tail impacts SSA repair and mismatch correction, PAM sequences were artificially inserted adjacent to each repeated fragment in FA Tailed strains so that cleavage would occur at one boundary of the repeats (Figure 2C). A Cas9-mediated DSB created adjacent to the left repeat results after annealing in a 2.3 kb nonhomologous tail (flap) on the bottom strand whereas a DSB created adjacent to the right fragment results in a 2.3 kb flap on the top strand. The viability in strains with only one flap was significantly lower compared to the Tailless strain (Figure 2D), suggesting that the presence of only one flap stimulates heteroduplex rejection. However, strains with only one flap had significantly increased viability compared to the FA Tailed strain in which repair requires the removal of two nonhomologous tails. The improved viability suggests that flap removal is a rate-limiting step in the process; however, it is possible that differences in the timing of cleavage of different Cas9::gRNAs could contribute to the fraction of colonies in which a repair event occurred. As discussed more below, the presence of colonies in which both F and A alleles are present at all sites could reflect a larger fraction of cells in which cleavage was only complete when cells had already replicated their genomes and thus could have two independent repair events.

### Effect of mismatch repair genes on SSA in Tailed strains

To confirm and extend our understanding of the roles of the MMR machinery and Sgs1 helicase in SSA repair (a possible mechanism illustrated in Figure S2), we deleted MMR genes and some helicases in the FA Tailed strain (Figure 3 & Table 1). As expected, *sgs4*Δ and *msh6*Δ increased SSA in the FA Tailed strain to nearly the level seen with AA, confirming that Sgs1 and Msh6 decrease SSA efficiency by promoting heteroduplex rejection [10, 26]. Deleting the Mph1 helicase did not show any effect on SSA repair (data not shown). Deleting *SRS2* resulted in very low viability in the FA Tailed strain (Table 1), consistent with previous studies that *srs2*Δ has a general defect in resolving recombination intermediates[5, 27, 28]. These results suggest that Sgs1 is the only 3’ to 5’ helicase required for heteroduplex rejection during SSA.

**Figure 3.**
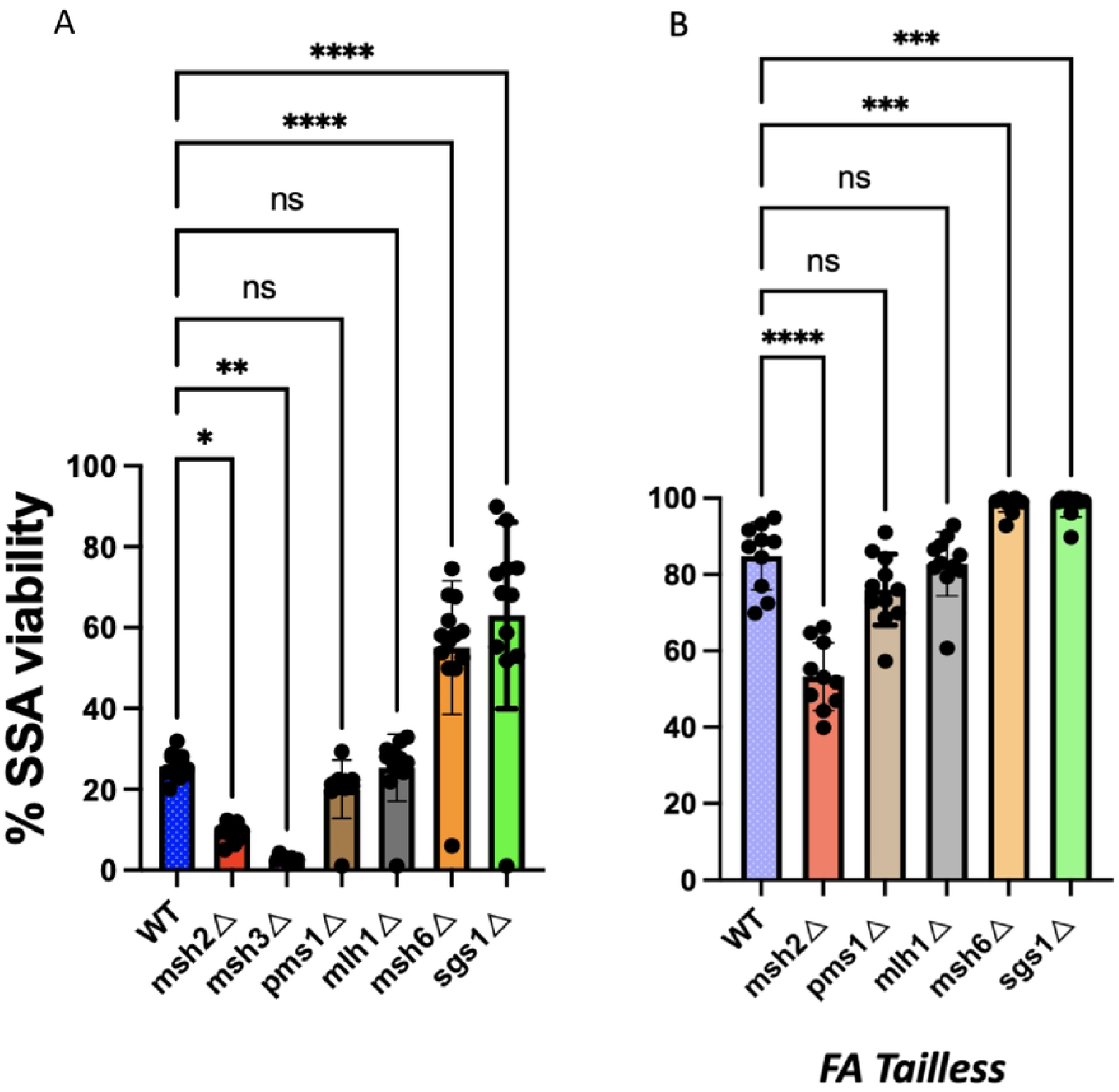
Viability of MMR and helicase mutants during SSA repair Effect of mismatch repair and helicase deletions on FA Tailed (A) and FA Tailless (B) strains. Experiments were independently repeated as described in Fig. 2A at least five times and averaged to arrive at the experimental mean. Means were statistically compared to WT Tailed or WT Tailless strains by One-Way ANOVA test. Error bars = SD. * p < 0.02; *** p = <0.001; **** p<0.0001.

Although Msh2-Msh6 has been shown to be involved in heteroduplex rejection and *msh6*Δ increased the repair frequency of the FA Tailed stain two-fold, *msh2*Δ resulted in a significant decrease in viability (10% recombination) (Figure 3). Since Msh2 also forms a complex with Msh3 (MutSß) and assists Rad1-Rad10 in removing nonhomologous tails [10], it is likely that the severe effect of *msh2*Δ compared to *msh6*Δ reflects its role in nonhomologous tail removal. Deleting *MSH3* also resulted in very low recombination (2.3%) in both FA and AA Tailed strains (Figure 3 & Table 1). Similarly, in *rad1*Δ Tailed strains the viability was only 1.5%. It is possible that the higher level of survival in *msh2*Δ (10%) versus *msh3*Δ or *rad1*Δ (2.3 or 1.5%) could be explained by the fact that *msh2*Δ - but not *msh3*Δ – could suppress heteroduplex rejection, thus increasing the number of survivors; however, while a *msh2Δ msh6*Δ double mutant yielded the same level of SSA as the *msh2*Δ mutant, a *msh3*Δ *msh6*Δ double mutant behaved as *msh3*Δ mutant, with only ~3% viability (Figure S3)-The deletion of MutL homologs (Mlh1 and Pms1) did not have a significant impact on viability in Tailed or in Tailless strains (Figure 3), confirming our previous results that the Mlh1-Pms1 endonuclease is not required for heteroduplex rejection in SSA [10].

### Effect of mismatch repair genes on SSA in Tailless strains

The high efficiency of repair observed in Tailless strains (Figure 2A) can be explained by there being no need for Msh2/Msh3-assisted tail removal by Rad1-Rad10. However, in the FA Tailless strain the viability decreased modestly to 85% compared to AA Tailless, suggesting that the recognition of MMs by the Msh2-Msh6 complex might have an inhibitory effect on SSA repair. Indeed, deleting *MSH6* increased SSA repair efficiency of the FA Tailless strain to almost the level seen for the AA Tailless construct. However, *msh2*Δ decreased survival to 53% (Figure 3 & Table 1), suggesting that there may be still another role for Msh2 in SSA involving divergent repeats. As expected, *sgs1*Δ also suppressed the effect of mismatches in the FA Tailless strain (Figure 3 & Table 1).

### The presence of nonhomologous 3’ tails affects the repair of mismatches in SSA

In strains with 3% divergence between the repeated fragments, the resected 3’ ssDNA tails will anneal to form a heteroduplex with six single-base mismatches and one “T” insertion distributed, as shown in Figure 1B. If these mismatches are not recognized and corrected by MMR proteins, they will exist as heteroduplex DNA and each strand will be used as a template for DNA replication so that both alleles will be found in SSA progeny as sectored colonies. If the MMs are corrected, then all descendants of the original cell will show the same genotype.

There is an ambiguity in the position of the insertion of a T (mismatch 2) within a run of 12 Ts. Because MM1 is also a T in in the F variant, there are actually 12 Ts in a row in F compared to 10 in the A variant. As we document below, there is a high level of discordance between the repair of MM2 compared to the adjacent MM1 and MM3, likely attributable to errors in PCR amplification and sequencing. Hence, in most of the analysis below, we have only reported the fate of the 6 well-behaved mismatches, omitting MM2. With strand linkage analysis (below) we could also specify MM2.

We investigated by Sanger sequencing how MMs are corrected during SSA repair (Figure 4). For the FA Tailed strain, where SSA was initiated by inducing HO endonuclease, approximately half of the outcomes (66/143) displayed correction of all mismatched sites either as all F or all A (Figure 4A); of these, 57 (86%) repaired all sites as F. Among the remaining 77 colonies there was a mixture of fully corrected F or A alleles as well as a minority with unrepaired heteroduplex. Overall, the correction of the six mismatches appears to display a gradient of correction such that sites near the left end are corrected as F whereas sites closer to the right end showed a higher proportion of corrections to A (Figure 4A). Very similar results were obtained when we directed cleavage at almost the same site by introducing a constitutive Cas9 plasmid pAB101 (Figure 4B). We note that MM7 is about 50 bp further from the end of the homology than MM1, so that we likely underestimate the bias of correction at the right side of the repeat.

**Figure 4.**
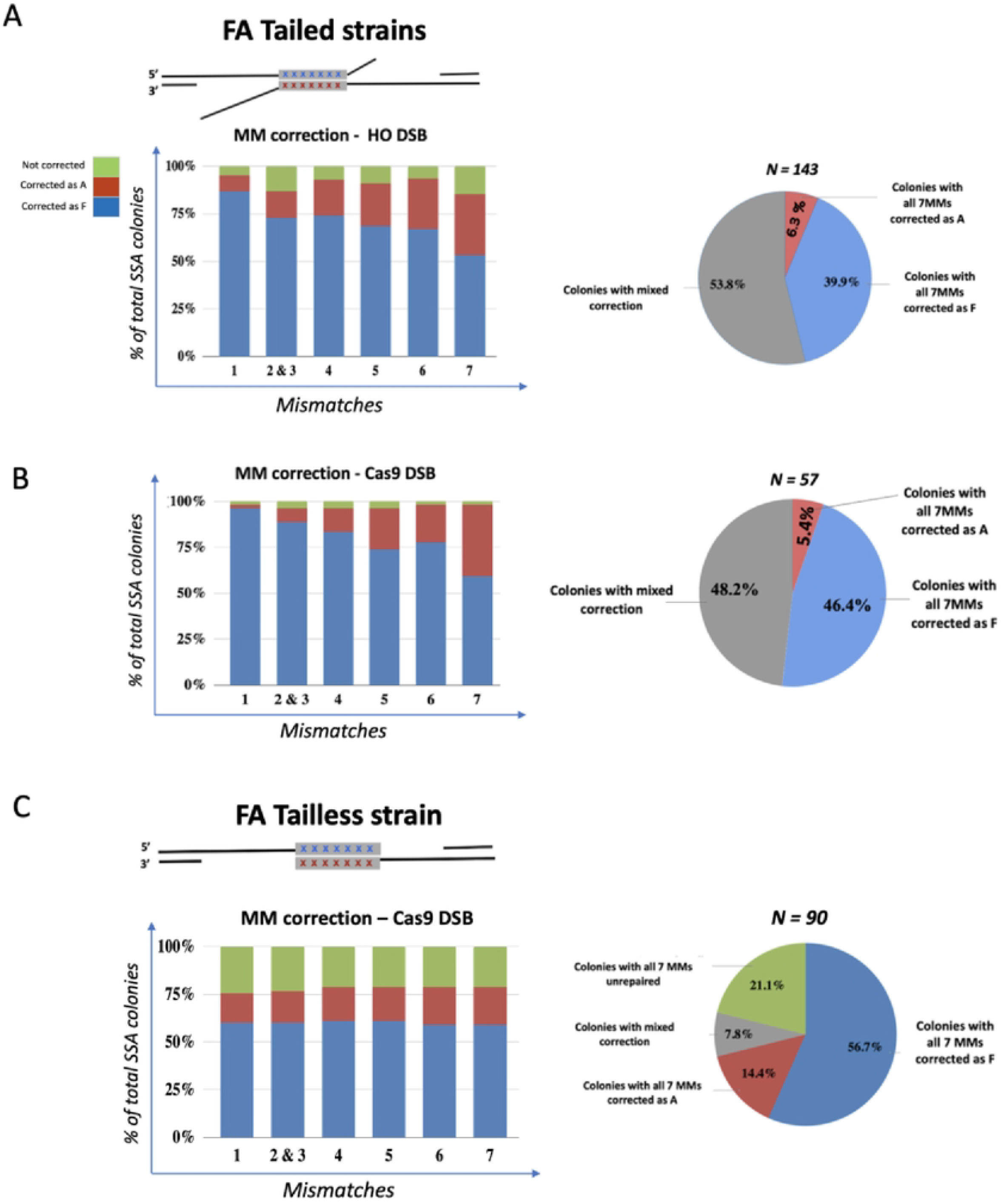
Sanger sequencing analysis: Mismatch correction in FA Tailed and Tailless strains. A. Gradient of mismatch correction across the SSA repeat in the FA tailed strain. Sanger sequencing analysis of 143 SSA repaired products resulted upon induction of *GAL::HO.* Left: the six columns from the chart correspond to each MM (from MM1 to MM7) except for MM2 and represent the percentage of survivals that repaired a mismatch as A (Red), as F (Blue) or remained unrepaired MM (green). Right: the pie shows the overall SSA repair outcomes. The majority of SSA colonies showed mixed repair (grey). In 40% of colonies (blue) all the mismatches were repaired as the F repeat and only 6% showed all the mismatches repaired as A (red) when DSB was created by HO. B. The chart (left) and the pie (right) show a similar gradient of correction in 57 SSA repaired colonies when a when a DSB is created by Cas9 (pAB101). C. Sequences for SSA products from FA Tailless strain showed uniform correction that favors the left fragment F fragment (blue).

We investigate the possibility that the indel located on the left side might cause the repair bias. We replaced the deletion of 1 T in the “A” sequence by inserting a G at position 23, 5 bp upstream of MM3 (Figure S4A & Supplementary Table 1), creating the strain designated as nFA. Again, there was a strong bias towards the repair of all alleles as F, with the highest level of F incorporation at the left end of the segment (Figure S4C).

The bias to correct sequences in favor of F is in fact a bias in favor of whichever allele is located close to the left end of the annealed structure. We created a Tailed SSA construct, designed as AF, in which the positions of the A and F segments are reversed (Figure S5A). When the A allele is located to the left of the DSB, repair now favors the A allele at the left end (Figure S5B), and, again, the replacement of the T deletion with a G to create a base-pair mismatch at MM2, 3bp from MM3 (strain AnF) yielded a similar pattern to that seen with the single base deletion (Figure S5).

In contrast, in FA Tailless strains, Sanger sequencing analysis of 90 colonies revealed a remarkably uniform pattern of correction of all the mismatches (Figure 4C). Similar to the FA Tailed strains, where 40% corrected all markers to F and only 6% corrected all to A, in the Tailless strain all 7 MMs were corrected in ~ 56% of events as F while ~15% were corrected to all A. But different from the FA Tailed strain, where the majority (54%) of survivors are the results of a mixed MM correction, only 8% of the SSA Tailless survivors (7/90 colonies) showed a mixed correction (Figure 4C&D). In 21% of the FA Tailless colonies, none of the 7 MMs was apparently repaired during SSA; these progeny presented a pattern where all of the sites contained both F and A alleles. The frequency of this outcome was lower in SSA repaired colonies from FA Tailed strain (only 10% according to stand linkage analysis, described below). One difference between the Tailed and Tailless strains is that SSA in the Tailed strain is induced by HO endonuclease cleavage, which occurs rapidly after galactose induction, whereas the Tailless strain employed a galactose-inducible Cas9 construct; moreover the site being cleaved is different and may be less accessible. It is possible that there is a delay in cleavage, such that a fraction of the colonies would have had independent DSB repair in the two daughter nuclei. In some of those cases one SSA event would yield only F alleles whereas in the second, only A alleles would be recovered, yielding what appeared to be persistent heteroduplex across the region.

Evidence that *GAL::CAS9* cleavage might be less efficient and possibly delayed comes from a similar galactose-inducible Cas9 cleavage of the Tailed strain, cutting in the middle of the phage λ sequences, directed by plasmid pES2, i.e., at a site different from that shown in Figure 4 (Figure S6A and B). We found that 4/26 (15%) colonies analyzed showed the presence of both F and A alleles at each site. Among the other outcomes, we saw the same strong bias in favor of complete correction of alleles to F (23%) compared to full correction to all A alleles (4%) that we found with HO-induced cleavage and the same gradient of correction favoring F alleles at the left end and A alleles towards the right end. These data reinforce the conclusion that mismatch correction in the Tailless strain is quite different from the Tailed strain, that the presence of the 3’ nonhomologous tail removal impacts the way mismatches are corrected in the SSA intermediate, and that this pattern can be seen even when Cas9-directed cleavage is less efficient than at other sites.

### Effect of mismatch machinery and helicase mutations on SSA repair

As previously described [10], *msh6*Δ, but not *pms1*Δ and *mlh1*Δ, increased the level of SSA in the Tailed FA strain compared to WT strain (Table 1 and Figure 3). Sequencing results, showed, as expected, all three mutations led to a strongly increased percentage of uncorrected MMs in the Tailed FA strain (Figure 5). This high percentage of heteroduplex was observed from MM3 to MM7 (Figure 5), suggesting that repair of these MMs is mostly dependent on mismatch machinery. However in *msh6*Δ, *mlh1*Δ and *pms1*Δ, MM1 still showed a significant percentage of mismatch correction. As this MM is located close to the extremity of repeated fragment (position +17 bp; Figure 1B & Table 3), it is possible that its correction, and perhaps also some of the corrections of MM2/3, can occur by an alternative mechanism (see below).

**Figure 5.**
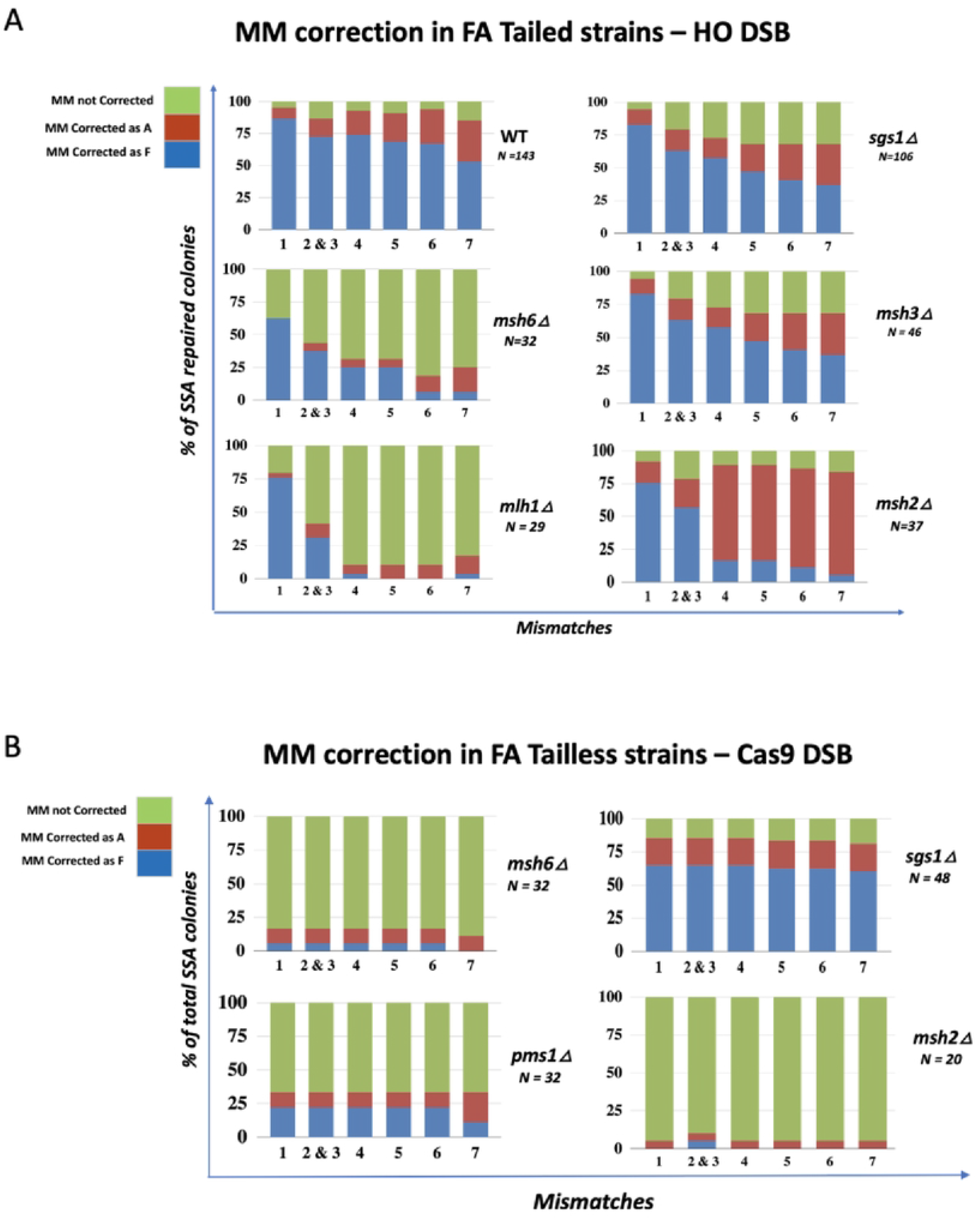
A. Impact of MMR machinery on MM correction during SSA in FA Tailed and Tailless strains. A. Mismatch correction in Tailed strains. Sanger sequencing analysis of SSA outcomes showing increased percentage of unrepaired MMs compared to WT in *msh3*Δ, *msh6*Δ, *mlh1*Δ, *and sgs1*Δ backgrounds. The *msh2*Δ strain showed an altered gradient that may result from a small population of survivors repairing by an alternative mechanism. Heteroduplexes (unrepaired MM) are shown in green. B. Uniform MM correction in mismatch repair mutants in FA Tailless strains.

As we noted above, Msh3 and Msh2 both play important roles in the clipping of 3’ nonhomologous tails; deletion of either gene reduced the efficiency of SSA to ≤10%. Among the low-frequency SSA events that occur in the absence of Msh2 the percentage of heteroduplex (unrepaired MMs) was very low (<20%) (Figure 5A). Moreover, the pattern of correction for *msh2*Δ was remarkably different than wild type for MM4, MM5, MM6 and MM7, as they were now repaired strongly in favor of the alleles on the bottom strand (A fragment). These data suggest that during SSA in the absence of Msh2 there is a small population of cells that might undergo repair through an alternative mechanism of nonhomologous tail removal. These SSA outcomes are characterized by a transition between between MM3 and MM4 that leaves MMs 1-3 mostly unrepaired while MMs 4-7 are usually repaired as A.

We then examined the roles of the mismatch repair genes and helicases when SSA occurred in the Tailless strain (Figure 5B). In the absence of nonhomologous tails, deleting *MSH6 and PMS1* led to the great majority of repair events retaining heteroduplex DNA at each marker, while *sgs1*Δ had little effect. Strikingly, however, in the FA Tailless strain, deleting *MSH2* yielded 70% fully sectored colonies (i.e. where no site was repaired), a result quite different from the outcomes in the Tailed strain where there was strongly directional repair of the all markers. In the Tailless strain, there is no requirement for 3’ tail clipping; consequently *msh2*Δ behaved quite similarly to *msh6* or *pms1Δ.* These data again suggest that the presence of nonhomologous tails affects mismatch correction. These data also demonstrate that there is a tail-dependent and Msh2-independent pathway, though inefficient, that leads to the recovery of SSA products with mismatch correction.

### Further characterization of mismatch correction by strand linkage analysis

To confirm the pattern of correction of the seven mismatches from heteroduplex DNA, we turned to Next Generation DNA sequencing of 96 independent SSA events as described in Materials and Methods. The analysis of all these NGS reads (see Materials and Methods) allowed us to characterize more completely the repair events occurring in each colony (see Figure 6). Strand-specific analysis showed which alleles were found in a given repair event and revealed which alleles were in tandem when some sites were uncorrected (Figures 6, S8 and S9). In most cases, MM2 was corrected as the adjacent MM3 and when MM3 was uncorrected (i.e. both F and A alleles were present in a single colony, MM2 was also uncorrected; but there is still a high level of discordance for MM2 that may reflect sequencing problems (Figure 6B and C).

**Figure 6.**
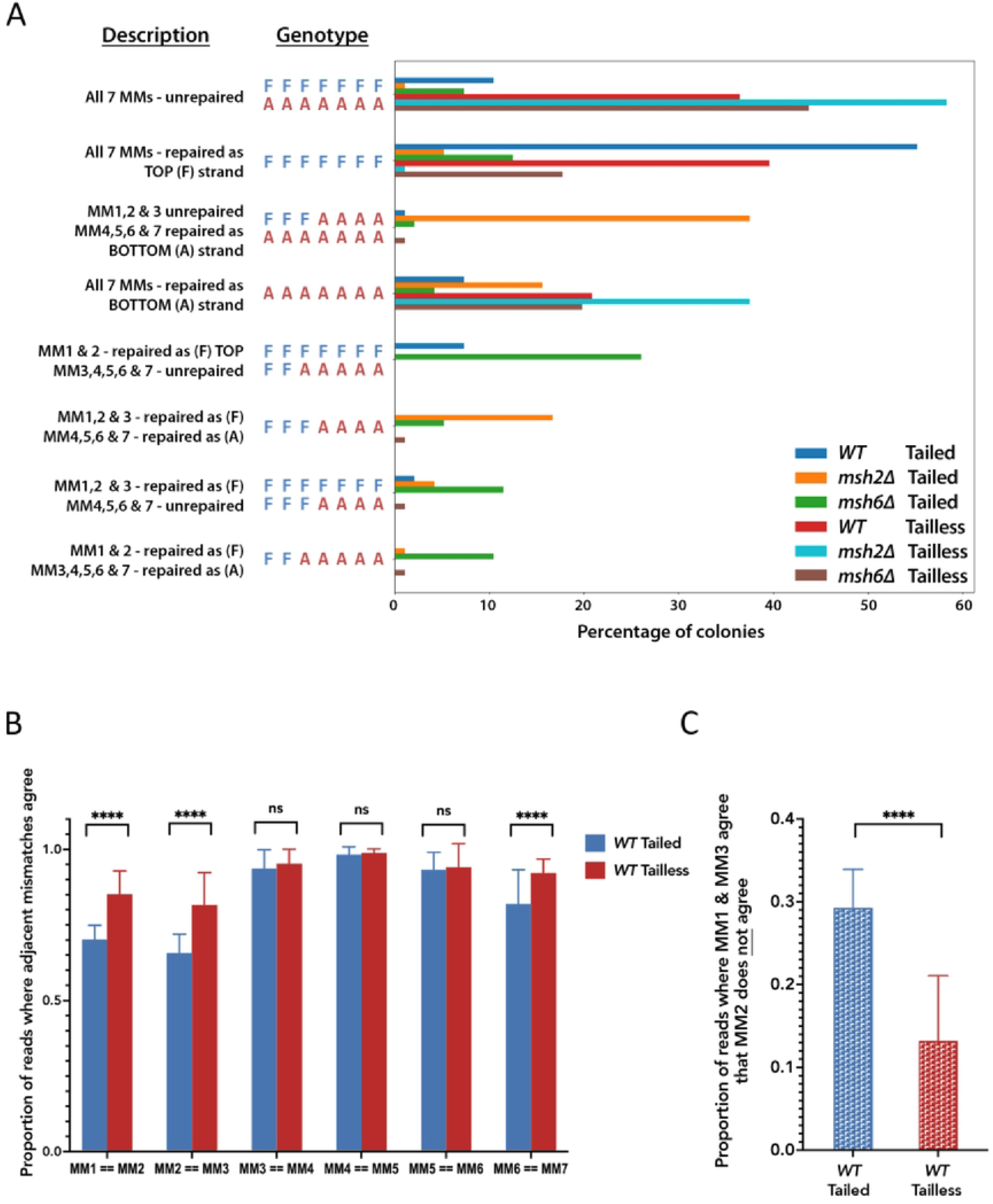
Representative genotypes of SSA repaired progeny analyzed by strand linkage analysis. A. Eight most frequent patterns of MM correction identified by analyzing all reads for each of 96 colonies per library and assigning them a genotype based on the most frequently observed read(s); see Material and Methods for details. Statistics for the number of reads per colony: mean ± sd = 186 ± 112; Interquartile range (IQR) = 111-241; Range = 7-1139. Only genotypes that appear more than 9 times for at least one of the strains are shown; complete results are shown in Figure S8. B. Concordance of mismatch correction with the next adjacent mismatch in Tailed and Tailless strains. The proportions (relative to the total reads) of reads containing alleles at adjacent mismatch positions that came from the same repeat. Comparisons are based on uncorrected data from the experiment shown in A. Statistical significance of differences among means were assessed by 2-way ANOVA. C. Non-concordance at MM2. The proportion of reads where MM2 did not match MM1 and MM3 are shown. Statistical significance of the difference between the means was assessed by an unpaired t-test. Values are means for 96 colonies per strain with error bars indicating standard deviations.. ns, non-significant (p>0.05); **** p<0.0001.

The most representative genotypes of six different libraries of SSA progeny, obtained from Tailed and Tailless FA WT, FA *msh2*Δ, and FA *msh6*Δ are shown in Figure 6A. In WT strains the most abundant pattern of correction showed a strong affinity for the F (left) repeat. Overall, this analysis shows that ~ 50% of the colonies from WT Tailed and ~ 40% from WT Tailless strains repaired all seven MMs in favor of the sequence of the F repeat.

Consistent with the results presented above, the highest percentage of sectored colonies (arising from post-repair replication of heteroduplex DNA) was observed in Tailless strains *msh2*Δ and *msh6*Δ (~ 60% and ~ 40%, respectively), where none of the 7 MMs was corrected/repaired. Some of these mixed outcomes likely represent failures of mismatch repair rather than any delay in cleavage. Unlike the Tailless strains, *msh2*Δ and *msh6*Δ Tailed strains resulted in an increased number of colonies with a mixed pattern of correction - where only some of the seven MMs were corrected. For example, in *msh6*Δ Tailed strain, the most representative genotype showed heteroduplex for mismatches MM3 through MM7 while the first two mismatches were repaired as the F alleles, again consistent with the results obtained by Sanger sequencing and suggesting that markers near the left end of the fragment may be repaired in a MMR-independent fashion. As we found with Sanger sequencing, the *MSH2* deletion in the FA Tailed strain produced a pattern of correction quite different from either WT or *msh6*Δ strains (Figure 6).

This strand-specific analysis also shows that in an *msh2*Δ Tailed background there is often partial MMR with the primary product being heteroduplex at the first 3 sites but fully corrected at the remaining 4 sites: FFFAAAA/AAAAAAA. Furthermore, the next two most abundant products are colonies with only FFFAAAA or only AAAAAAA. It is possible that a cell with the partially corrected heteroduplex arrangement (FFFAAAA/AAAAAAA) then replicated, but only one of the two daughters survived; this might occur if one of the two strands of the SSA product did not complete tail processing (as Msh2 plays an important, but not essential, role in that process) and only one strand was ligated prior to replication. Alternatively, there could be more than one round of mismatch correction before cells replicated.

While the *msh6*Δ Tailed strain produces many different products, the primary one (FFFFFFF/FFAAAAA) is the inverse of the *msh2*Δ, with FFFFFFF/FFFAAAA also being abundant. Moreover, similar to Tailed *msh2*Δ, the next most abundant outcomes appear to be the result of either another round of mismatch repair before replication or failure to recover both replication products, yielding only FFFFFFF, FFAAAAA or FFFAAAA.

In addition, this analysis reveals that even the WT Tailed strain produces a heterogeneous pool of products with about 10% showing a partial repair of one strand and more than 10% having no evidence of repair.

## Discussion

Single strand annealing (SSA) repair in budding yeast is a Rad52-dependent, but Rad51-independent repair pathway, occurring when a DSB is created between homologous sequences as short as 12-15 bp but optimally ≥ 200 bp [29, 30]. Here we show that the presence of nonhomologous DNA between two divergent direct repeats, leading to single-stranded 3’-ended nonhomologous tails (3’ flaps), affects not only the efficiency of recombination through SSA repair, but also the DNA sequence of the repair products.

In our system, the seven mismatches within the SSA intermediate are known to be recognized by mismatch proteins Msh2-Msh6 [13, 31, 32]. Once Msh2-Msh6 binds to the mis-paired nucleotides, the complex apparently recruits Sgs1-Rmi1-Top3 to stimulate heteroduplex rejection (Figure S2). However, even in AA Tailed strains, where the repeated fragments are identical, there was a significant increase of viability from 68% to 100% when Sgs1 was deleted (Table 1). This finding suggests that recruitment of Sgs1 and its associated proteins to heteroduplex DNA can be independent of the presence of mismatches themselves and may reflect recruitment of Sgs1 by the Msh2-Msh3 complex that recognizes the branched structure created by the presence of the 3’ nonhomologous tail (Figure S2). It is possible that even in the absence of mismatches, the inherent possibilities of misaligning the region of 12 Ts could provoke the attention of Msh2-Msh6.

Whether the timing of nonhomologous tail recognition and removal influences the correction of the mis-paired bases is not known. The processing of the two flaps may not be accomplished at the same time, perhaps depending on the DNA sequence identity at the branched structure. Apparently, SSA with only one nonhomologous tail is more efficient than substrates with two flaps (Figure 2C & D); but we cannot rule out differences in the efficiency of Cas9 cleavage where delayed cleavage would allow each daughter cell to repair independently and show an apparent increase in viability.

### 3’ flap structure dictates the bias of mismatch repair

The data presented here, along with previous studies [10] show that Msh2-Msh6, but not Pms1-Mlh1, are required for heteroduplex rejection; however, the correction of the mismatches themselves requires both Pms1 and Mlh1. Our data suggest that the decision to correct a given mismatch in favor of one allele or the other is strongly influenced by the presence of a 3’ nonhomologous flap, which favors the retention of the sequence on the 5’ end of the strand opposite of the flap. It is unclear when mismatch correction occurs relative to the Msh2-Msh3 aided Rad1-Rad10 mediated flap removal, but heteroduplex rejection most likely occurs before 3’ tail clipping. We suggest that flap cleavage precedes mismatch repair and that the newly-created 3’ end after flap removal may serve as the guide to repair mismatches favoring the opposite strand (Figure 7). Possibly the cleaved end would be perceived in the way that a nicked strand is designated to be the corrected strand during DNA replication [33]. The presence of a nearby 3’ end can also employ the 3’ to 5’ exonuclease activity of DNA polymerase δ to chew back as far as 25 nt before then polymerizing new DNA (Figure 7C), similar to strongly biased repair of an invading strand during homologous recombination [14, 34, 35]. The processing of the 3’ end in this way would suggest that some of the gradient of repair that we observed is attributable to this alternative process (Figure 7). We note that there is no such increase in the correction of MM1 in the tailless strain, implying that the processing of the tail itself may play a role in triggering this second mode of correction.

**Figure 7.**
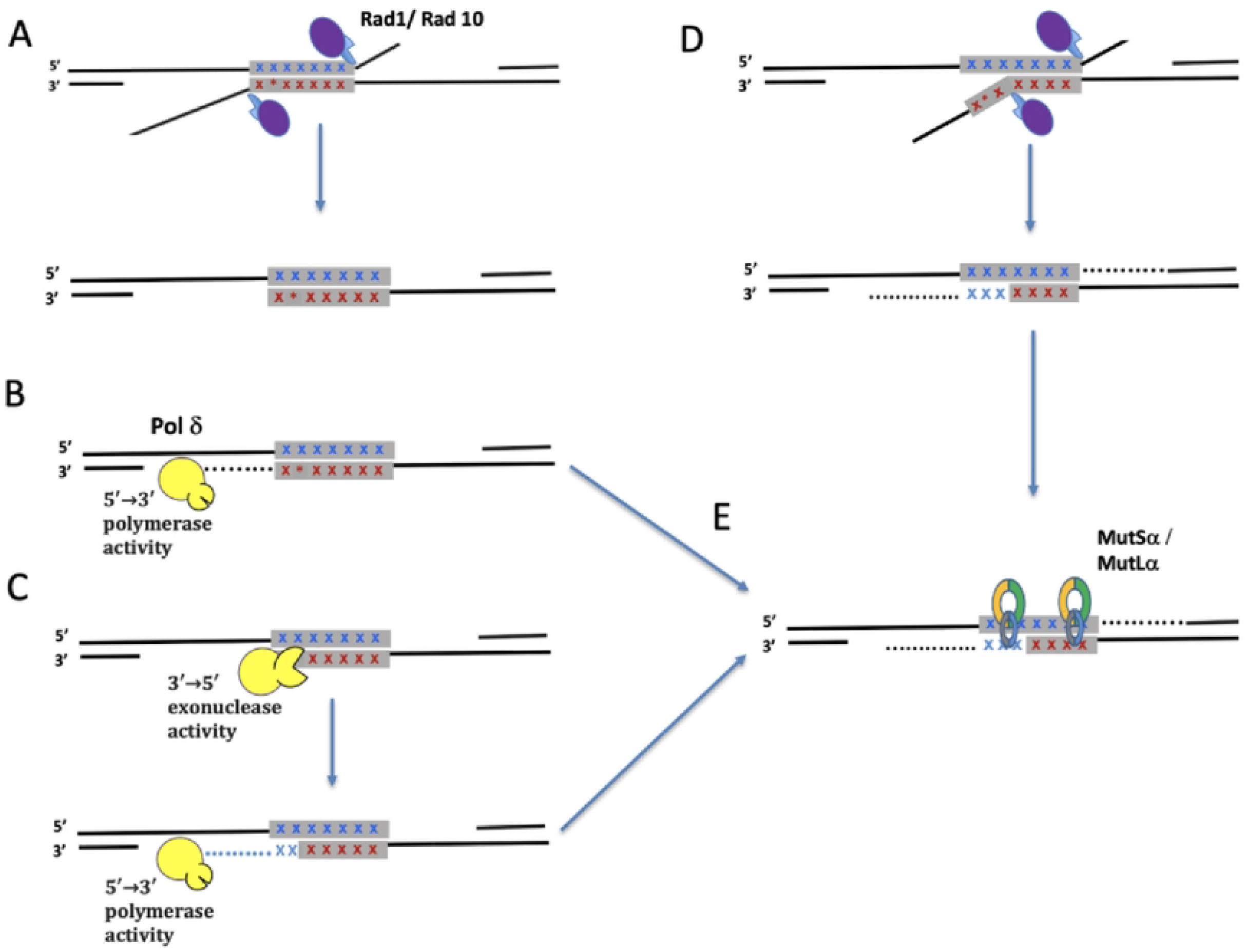
Pathways leading to mismatch correction of heteroduplex DNA formed during SSA. Heteroduplex DNA between the F and A variants has six single base-pair differences (X) and an insertion (*) that lies in a run of 12 Ts. A. Strand annealing is followed by Rad1-Rad10 clipping of 3’-ended nonhomologous tails. B. The 3’ ends can then be used as primers for Polδ DNA polymerase to fill in the gaps but C. in some cases Polδ’s 3’ to 5’ proofreading exonuclease activity may be stimulated to chew back the 3’ end and remove heterologies as far as 25 nt from the end, including the most terminal mismatches. D. The mismatches close to the 3’ end might also be excised by Rad1-Rad10 if the 3’ end is not firmly paired to its complementary strand. E. Mismatch repair performed by MutSα and MutLα is likely biased to remove mismatches on the filled-in (and perhaps not yet ligated) strand, creating a gradient of repair across the 200-bp region.

Recently the influence of the length of the 3’ nonhomologous flaps in directing mismatch repair during SSA in mammalian cells has been addressed. In a model with a single mismatch in a 152-bp segment, SSA events exhibited a modest 60-80% bias in correcting the mismatch prompted by the shorter of the two nonhomologous tails (33 nt vs. 11 nt) and that the pattern is reversed when cleavage creates tails of 0 and 42 nt [36]. Another recent study by Trost et al. [37] found that mismatches proximal to a nonhomologous tail were preferentially removed. However, unlike in budding yeast, Mlh1 as well as Msh6 caused a reduction in SSA outcomes with mismatched substrates. In yeast, Mlh1 and Pms1 are needed for mismatch correction per se but showed no effect on heteroduplex rejection. Trost et al. also examined the bias in repair when one nonhomologous tail was 268 nt long and the other varied from 16 nt to 9100 nt. Longer tails impaired SSA both for perfectly matched 267-bp repeats or for 1-3% divergent sequences. A marked gradient in repair, favoring retention of the allele on the strand opposite the nonhomologous tail, proved to be independent of varying the length of the second nonhomologous tail.

In our system, 3’ tails are hundreds of nucleotides long but the pattern of repair was not reversed when we used Cas9 to create cases where one end or the other lacked a tail. Instead, there seems to be some intrinsic bias in the region that strongly selects the left repeat (whether F or A) to be used as the template for correction. This bias is especially evident for MM1-MM3. It is possible that the presence of both an indel at MM2 and the mismatch at MM3 could create a mismatched site that is recognized more strongly than other sites and influences the repair outcome; however, when we replaced the indel with a T:G mismatch, separated by 5 bp from MM3, the overall spectrum of mismatch correction was unchanged (Figures S4 and S5).

The strong bias for retaining the sequence of the top strand near the left end of the segment was also observed when creating a DSB adjacent to left fragment leading to one nonhomologous tail, localized 17bp upstream MM1 (Figure S7A). These results likely reflect a second mechanism of correction involving DNA polymerase δ (Figure 7). We have recently showed that the proofreading 3’ to 5’ activity of DNA polymerase δ can resect as far as 40 nt from the 3’ end after Rad51-mediated strand invasion [10, 14, 34]. A similar 3’-5’ removal of the 3’ end after flap removal will also favor incorporation of the sequence on the top strand (Figure 7). When a DSB was created adjacent right fragment, leading to one nonhomologous tail on the top strand, the most proximal mismatch, MM7 (localized 70bp downstream the 3’ tail), was less likely to be corrected (Figure S7B). This suggests that correction of the MMs localized in the middle of the SSA intermediate depend on MMR machinery. The increased percentage of heteroduplex for MM7 (from 10% - in Figure S7A compared to 50% - in Figure S7B) can be explained by lower accessibility of MSH2/MSH6 to bind the mispaired nucleotides from MM7 while MSH2/MSH3 complex is engaged in nonhomologous tail removal.

The pattern of mismatch repair suggests that in Tailed strains there are often partial repair events that terminate (or start) near MM3 resulting in “patchy” mismatch incorporation. However, in Tailless strains, the assimilation of each mismatch occurred at the same frequency at each location. We note also that there is a bias in favor of retaining the sequences of the left repeat, whether it is F or A. This result suggests that sequences adjacent to the repeats exert some effect on the repair process; but at this time we do not know what DNA sequence features, alterations in chromatin structure or effects of nearby transcription might be. The repair often seems to be “all or none” across at least the 2/3 of the sequence that is marked. These results imply that there is a broad excision/repair event that covers most of the region. Whether mismatch repair happens before the DNA is filled in and ligated is not known. In both Tailed and Tailless strains, Msh2 was found to play a central role, as this protein, depending on its partners, is involved in critical steps during SSA repair: heteroduplex rejection, nonhomologous tail removal and mismatch correction. We note that in contrast to wildtype and *msh6*Δ mutants, the viability of *msh2*Δ Tailed strains is very low; moreover, the pattern of mismatch correction is almost the opposite of what occurs in *msh6Δ.* We suspect that there is another, low-efficiency pathway that can operate without Msh2 that would account for these unusual outcomes.

## Acknowledgements

We are grateful for comments on the manuscript from Jeremy Stark, Michael Lichten, Sue-Jinks-Robertson and members of the Haber lab. Rebecca Tsai helped in the construction of the tailless strain. This work was supported by NIH grant R35 GM127029.

